# Circumferential actomyosin bundles drive endothelial cell deformations to constrict blood vessels

**DOI:** 10.1101/2024.12.22.630001

**Authors:** Yan Chen, Nuria Taberner, Jason da Silva, Igor Kondrychyn, Nitish Aswani, Guihua Chen, Yasushi Okada, Anne Karine Lagendijk, Satoru Okuda, Li-Kun Phng

## Abstract

Following the formation of new blood vessels, vascular remodelling ensues to generate a hierarchical network of vascular tubes with optimal connections and diameters for efficient blood perfusion of tissues. How transitions in endothelial cell (EC) number and shape are coordinated to define vessel diameter during development remains an open question. In this study, we discovered EC deformations, rearrangements and transient formation of self-seam junctions as key mechanisms that explain a negative relationship between cell number and vessel diameter. High-resolution analysis of actin cytoskeleton organization disclosed the generation of tension-bearing, circumferential actomyosin bundles in the endothelial cortex that drive EC deformation and vessel constriction. Importantly, the loss of circumferential actin bundles in *krit1*/*ccm1*-deficient ECs causes cell enlargement and impaired vessel constriction that culminate in dilated vessels, characteristic of cerebral cavernous malformation. Our multiscale study therefore underpins circumferential actomyosin-driven EC deformations in controlling vessel size and in the prevention of vascular malformations.

## Introduction

Tissue growth and function critically depends on the efficient distribution of oxygen and nutrients through a hierarchical network of larger arteries and veins and smaller capillaries of optimal connections. While directed endothelial cell (EC) migration initially shapes the pattern of the primitive vascular network, its final organization is determined by vessel remodelling, which comprises of vessel pruning and diameter regulation.

The formation of correctly sized blood vessels is essential for tight control and efficient distribution of blood flow since lumen diameter significantly influences flow resistance^1,2^. The mis-regulation of vessel size can culminate in vascular malformations such as Hereditary Haemorrhagic Telangiectasia (HHT), which is characterized by enlarged arteries and veins that connect with each other to create shunts that siphon blood from capillaries, and Cerebral Cavernous Malformation (CCM), which presents as clusters of abnormally dilated blood vessels in the brain^3–5^. In many vascular malformations, blood flow distribution through the vascular network is impaired and can result in bleeding and chronic pain^6^. Furthermore, outward and inward remodelling of blood vessels in response to changes in heart rate and cardiac output is an essential mechanism in maintaining a shear stress set point and tissue homeostasis^7^. The elevation in blood flow causes vessel expansion that decreases shear stress while the reduction in flow results in the narrowing of vessels to increase vascular resistance and shear stress. Given the importance of vessel size in health and disease, it is imperative to uncover cellular mechanisms of vessel diameter regulation to understand the aetiology of vascular malformations, many of which form during early vascular development, as well as in the regulation of tissue homeostasis.

Previous studies have demonstrated that vessel diameter is regulated autonomously by EC number and cell size. In the mouse yolk sac, the fusion of separate vessels into a single vessel leads to an increase in EC number and results in a dilation of the vessel^8^. When directed migration of ECs within the vascular network is perturbed and causes misdistribution and abnormal accumulation of ECs, vessels enlarge^9–11^. Additionally, in the adult mouse brain, the diameter of capillaries is positively correlated with EC number^1^. However, other studies demonstrate that vessel diameter is controlled by cell size independent of cell number^12,13^. Blood flow-induced dilation of blood vessels requires an increase in EC size that is dependent on PlexinD1-KLF2 mechanosensitive signalling^14^. The Bone Morphogenetic Protein (BMP) pathway also regulates vessel diameter by controlling EC size. The depletion of Alk1, a transforming growth factor beta (TGFβ) receptor, its coreceptor Endoglin^12,15^, or the downstream mediator, SMAD4^16^, results in enlarged ECs that lead to aberrations in vessel diameter as observed in HHT. EC size is also increased in models of CCM with KRIT1/CCM1 mutation^17–19^. How these signalling pathways regulate EC size is still unclear. As recent studies have implicated a role of actin cytoskeletal remodelling in controlling EC size and vessel diameter^13,20,21^, it is likely that alterations in EC mechanics may underlie the vascular anomalies.

In this study, we have investigated the role of EC behaviours and mechanics in regulating the remodelling of blood capillaries using zebrafish intersegmental vessels (ISVs), which undergo constriction and elongation from 2 days post fertilisation (dpf). By performing high-resolution timelapse imaging and cell and tissue strain analysis, we discovered that the remodelling of ISVs is driven by the combined activities of EC deformation and changes in cell number arising from cell divisions and cell rearrangement. At the subcellular level, we identified dynamic and distinct actin organizations in the EC cortex. Crucially, we observed that tension-bearing circumferential actin bundles extend from cell junctions and/or plasma membranes to the cortex and that their formation coincides with vessel constriction. The disruption of circumferential actin formation or inhibition of myosin II activity increases vessel diameter, pointing towards a function of circumferential actomyosin-driven EC deformation in vessel constriction. To examine the importance of this mechanism in regulating vessel constriction, we analysed ECs deficient in Krit1. *krit1*-deficient ECs lack circumferential actin, undergo reduced EC deformation and exhibit enlarged vessels. These findings therefore implicate an important role of circumferential actin in driving EC deformations to generate vessels of correct size.

## Results

### Cell deformation, cell rearrangements and changes in cell number underly vessel remodelling

To investigate the temporal changes in vessel morphometrics during vascular remodelling, we examined zebrafish arterial and venous intersegmental vessels (aISVs and vISVs) from 2 to 4 dpf (Fig. 1a). Both vessel types exhibit a significant reduction in diameter and an increase in length over this period, indicating vessel constriction (Fig. 1b) and elongation (Fig. 1c), respectively. As EC number contributes to vessel size^8,9^, we examined how EC number changes during vascular remodelling. Quantification of EC nuclei revealed a substantial increase in cell number within both aISVs and vISVs during this period (Fig. 1d), demonstrating an inverse relationship between cell number and vessel constriction, and a positive correlation between cell number and vessel elongation.

**Figure 1.**
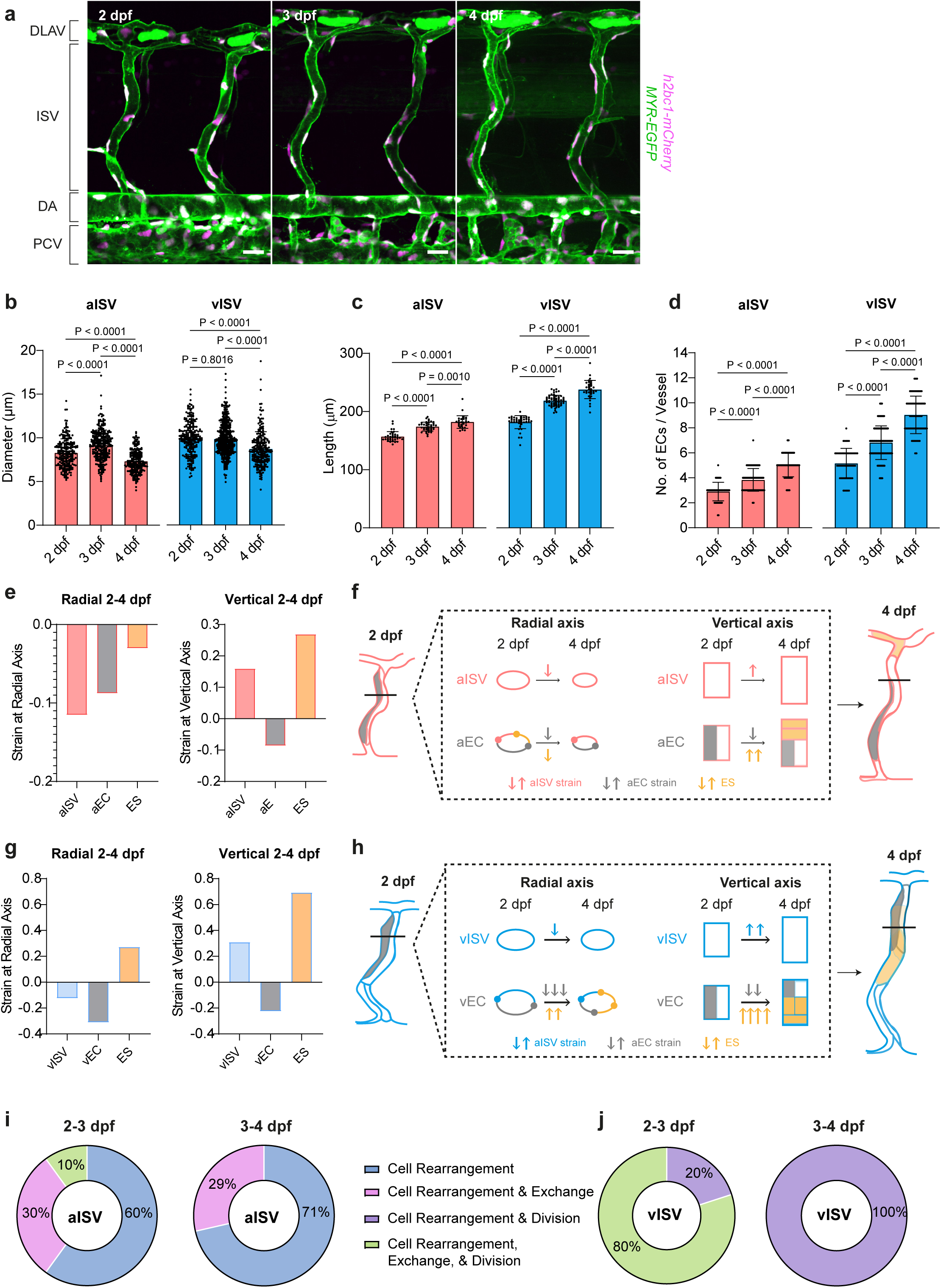
Changes in cell size and cell number contribute to vessel constriction and elongation during vessel remodelling. **a** Maximum intensity projection of confocal z-stacks of ISVs in zebrafish trunk from 2 dpf to 4 dpf. *Tg(fli1:h2bc1-mCherry)^ncv^*^31^ and *Tg(fli1:MYR-EGFP)^ncv^*^2^ label endothelial cell nuclei and cytosol, respectively. Scale bar, 20 µm. **b** Quantification of vessel diameter from 2dpf to 4dpf in aISV and vISV (n=228/284/213 aISVs and n=228/500/240 vISVs at 2/3/4 dpf). **c** Quantification of vessel length from 2dpf to 4dpf in aISV and vISV (n=36/40/30 aISVs and n=40/60/34 vISVs at 2/3/4 dpf). **d** Quantification of number of ECs from 2dpf to 4dpf in aISV and vISV (n=57/71/54 aISVs and n=52/127/65 vISVs at 2/3/4 dpf). Statistical significance was assessed by ordinary one-way ANOVA with Turkey’s multiple comparisons test. Mean values are indicated (**b-d**). **e** Strain analysis at radial and vertical axis in aISVs (average from **b**, **c**), aECs (average from n=16/32 cells at 2/4 dpf), and ES of aECs between 2 dpf and 4 dpf (see source data). **f** Schematic of strain analysis to show the changes in aISV and aEC between 2 dpf and 4 dpf during vessel constriction and elongation. **g** Strain analysis at radial and vertical axis in vISVs (average from **b**, **c**), vECs (average from n=13/21 cells at 2/4 dpf), and ES of vECs between 2 dpf and 4 dpf (see source data). **h** Schematic of strain analysis to show the changes in vISV and vEC between 2 dpf and 4 dpf during vessel constriction and elongation. **i,j** Percentage of ECs undergoing cell rearrangement, exchange, division or combinations of these events in aISVs (2-3 dpf: n=10; 3-4 dpf n=7) and vISVs (2-3 dpf: n=5; 3-4 dpf n=6). DA dorsal aorta; DLAV dorsal longitudinal anastomotic vessel; PCV posterior cardinal vein; ISV intersegmental vessel; aISV arterial ISV; vISV venous ISV. Source data are provided as a Source data file.

Although the increase in EC number can account for the elongation of aISVs and vISVs, it does not explain the decrease in their diameter during vascular remodelling. We therefore sought to clarify the contribution of EC shape changes to vessel constriction by analysing cell size and cell aspect ratio using a custom ImageJ script^13^. We found that venous ECs (vECs) showed a significant reduction in cell size without a corresponding change in aspect ratio, while arterial ECs (aECs) showed no significant changes in either parameter (Supplementary Fig. 1a & b). This might be because the diameter (∼10 μm) is much smaller compared to the length (∼100 μm), so any change in diameter is cancelled out by the length when calculating the area. For example, even though the diameter and length of ISVs changed considerably, the calculated vessel area (based on these two dimensions) stayed consistent between 2 and 4 dpf, particularly for aISVs (Supplementary Fig. 1c). This stability suggests that measuring cell area may not be a sensitive and reliable measure to detect changes in cell dimensions and could explain why significant size changes were not observed. To address this, we applied strain analysis, defined as the ratio of dimensional changes between 2 and 4 dpf. Tissue strain quantifies overall vessel deformation, while cell strain measures changes in individual EC width and height, reflecting direct cell shape deformation. To assess cell configuration changes, which is driven by cell division and cell rearrangement, we also calculated effective strain due to cell number change (hereafter referred to as ES). This parameter represents changes in the effective number of cells required to occupy the vessel’s perimeter or length based on EC width or height. By comparing cell strain and ES, this analysis distinguishes two key contributions to vessel deformation: 1) cell deformation, quantified by cell strain, which reflects changes in cell shape; 2) cell configuration changes, quantified by ES, which capture the effects of cell division and rearrangement. This approach clarifies the distinct roles of cell shape and configuration changes in driving vessel remodelling.

In aISVs, radial tissue strain is negative (Fig. 1e), consistent with the observed reduction in vessel diameter (Fig. 1b). Similarly, radial cell strain is also negative, reflecting a reduction in cell width. In contrast, vertical tissue strain is positive, indicating vessel elongation, while vertical cell strain is negative, reflecting reduced cell height. This resulted in a positive ES along the vertical axis (Fig. 1e). These observations suggest that cells reduce in height along the vertical axis as the vessel elongates, with elongation driven by an increase in cell number. However, ES is negative at the radial axis (Fig. 1e). The inverse trends in ES along the two axes suggest cell rearrangement, where cells move along vertically, thereby reducing cell width or cell number along the radial axis while increasing it vertically (Fig. 1f)

In vISVs, radial tissue strain is negative and vertical tissue strain is positive, consistent with vessel constriction and elongation between 2 and 4 dpf. Meanwhile, cell strain is negative along both axes (Fig. 1g) suggesting significant reduction in cell size. As a result, the ES is positive along both axes, suggesting that considerable contraction of vECs along both axes is important to accommodate the expanded cell population within the vISVs (Fig. 1h).

In summary, results from our morphometric analyses implicate EC deformation and cell rearrangements as key cellular behaviours that govern vessel remodelling, while increased cell number promotes vessel elongation.

### EC proliferation and migration differentially regulate cell numbers in aISVs and vISVs

To understand the coordination between cell division and deformation, we performed time-lapse imaging to observe the dynamic behaviours of ECs in the endothelial-specific reporter line *Tg(fli1:GAL4FF)^ubs^*^3^*; Tg(UAS:EGFP-UCHD)^ubs^*^18^, which labels both cortical and junctional filamentous actin (F-actin). Imaging was performed from 2 to 3 dpf and 3 to 4 dpf. To assess changes in EC number, we tracked EC divisions as well as cell migration by recording the number of ECs entering or exiting the ISVs (cell exchange, supplementary Fig. 2a). Between 2 and 3 dpf, 30% of aISVs exhibit cell exchange, while 10% display both cell division and exchange (Fig. 1i). Importantly, all cell exchanges occur exclusively between aISVs and the dorsal longitudinal anastomotic vessel (DLAV), with none observed between aISVs and the dorsal aorta (DA) (Supplementary Fig. 2b), consistent with prior findings showing increased EC numbers in aISVs at 1 dpf due to immigration from the DLAV, not the DA^10^. While the earlier study reported 60% of aISVs undergoing cell exchange at 1 dpf, our findings reveal a decline at later stages (2 – 4 dpf), with most aISVs (60%-70%) exhibiting only positional rearrangements (Fig. 1i).

At 2-3 dpf, 80% of vISVs exhibit both cell division and exchange (Fig. 1J). More cells contribute to the DLAV than are received from the posterior cardinal vein (PCV) (Supplementary Fig. 2c), suggesting that the net increase in EC number in the vISVs is primarily due to cell division. All observed vISVs undergo mitosis, 40% experiencing a single division event, and 60% undergo two or three (Supplementary Fig. 2d). By 3 – 4 dpf, vISVs cease cell exchange and exhibit only division, with half undergoing one division event and the other half experiencing two (Supplementary Fig. 2e).

Collectively, these results indicate that the increase in EC number in aISVs results mainly from EC exchange, whereas in vISVs, it is largely due to cell division.

### Junction remodelling and cell deformation contribute to vessel constriction in ISVs

While ISVs that undergo cell exchange with the DLAV naturally exhibit active cell migration, we also observed highly dynamic cell rearrangements within ISVs (Fig. 1i & j). To monitor the dynamic deformation and rearrangement of cell-cell junctions in ISVs, we visualized actin filaments using EGFP-UCHD at the front and rear of the vessel (Fig. 2a and c).

**Figure 2.**
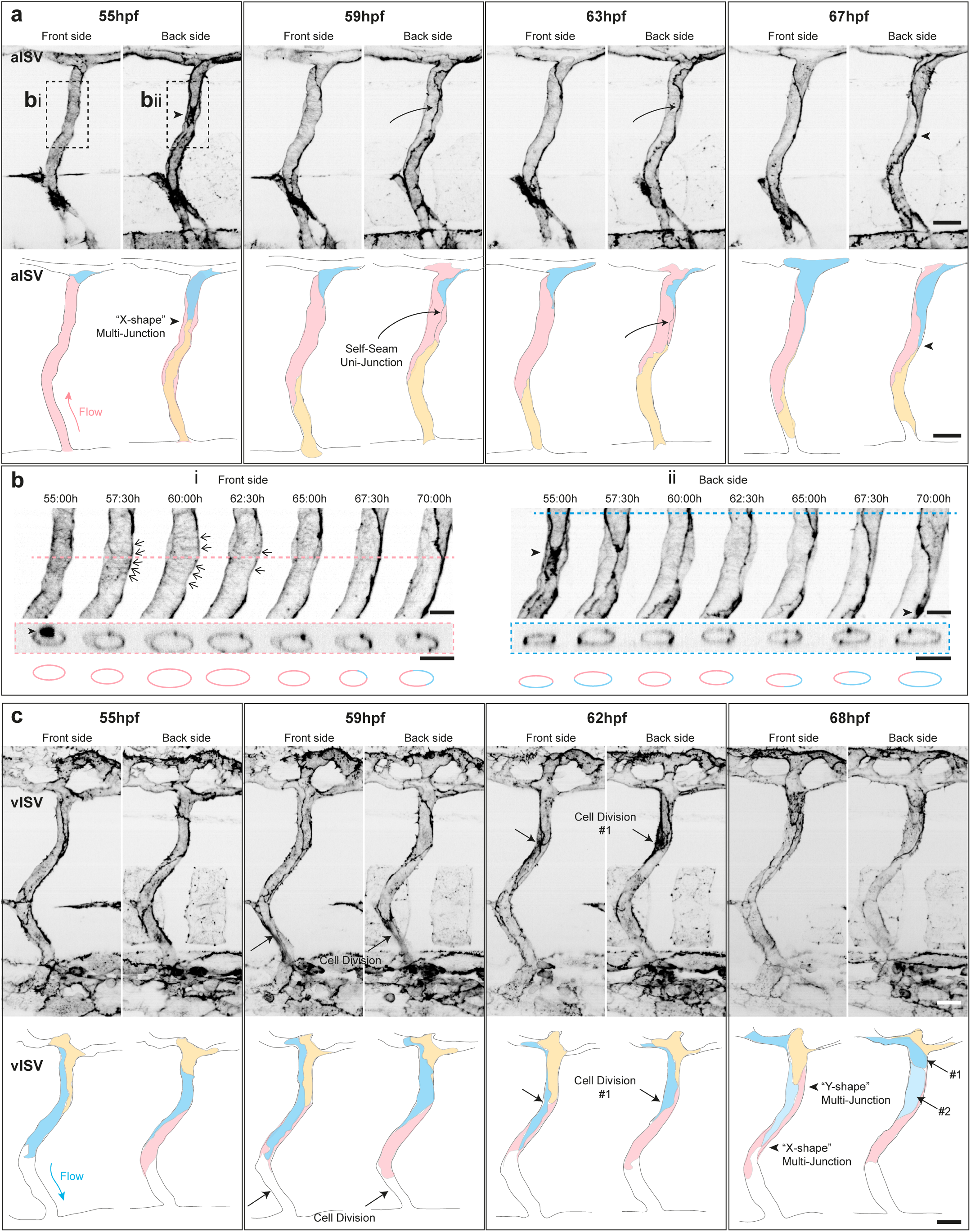
Distinct dynamics of cell rearrangement and cell deformation during vessel remodelling. **a-c** Still images of an aISV (**a, b,** Supplementary Movie 1) or a vISV (**c,** Supplementary Movie 2) showing cortical and junctional actin in embryos from *Tg(fli1:GAL4FF)^ubs^*^3^*; Tg(UAS:EGFP-UCHD)^ubs^*^18^ at 2-3 dpf. The blood vessel is separated as the front side (left channel) and the back side (right channel). Schematics are traced from the original image and denote the constituent cells in colour (pink, blue, and yellow). Multi-junction (black arrowhead) indicates junction that is formed by two or more cells, while Uni-junction (black arrow) indicates junction that is formed by a single cell. Similar observations were made in 10 movies. Scale bar, 20 μm. **b** Magnification of the inset in **a**. Cross-section showed at z-plane indicated by pink serrated line (**i**) or blue serrated line (**ii**). Scale bar, 10 μm. **c** Cell divisions are observed and indicated by black arrows. Daughter cells (#1 and #2) are denoted in dark and light blue. Similar observations were made in 5 movies. Scale bar, 20 μm. ISV intersegmental vessel; aISV arterial ISV; vISV venous ISV.

In aISVs, blood flows in the direction from the ventral to the dorsal part of the vessel. Cells are seen migrating against the flow and moving from the DLAV to enter the aISV (Fig.2a, cell labelled in blue). Another cell (labelled in pink) established self-contact as well as intercellular adhesions with neighbouring cells (Fig.2a, labelled in blue and yellow), leading to the formation of an “X-shaped” multicellular junction involving three cells (Fig. 2a & bii, black arrowheads, and Supplementary Movie 1). This self-contact extends to form a transient self-seam junction, temporarily maintaining the vessel region as a unicellular tube (Fig. 2a, 59 hpf, black arrow and pink cell). Cross-sectional analysis confirmed this, showing that the single cell exhibited size oscillations, which gradually diminished over time (Fig. 2bi). During this period, stripes of circumferential actin were observed in the cell cortex (Fig. 2bi, black arrows). The self-seam junction persisted for approximately 10 hours, gradually shortening while maintaining cell-cell adhesions with neighbouring cells. At 67 hpf, the three cells re-established the “X-shaped” multicellular junction as they converged again. By the end of the timelapse recording, the cell in blue predominantly occupied the dorsal region of the vessel, contributing to the elongation of the vessel. The formation of self-seam junctions and associated cell rearrangements are consistently observed across all examined aISVs (n = 10/10), suggesting a stereotypical pattern of cell behaviour in aISVs starting at 2 dpf. Following these rearrangements, specific regions of the aISV (dorsal, middle, or ventral) transitioned from being shared by multiple cells to being predominantly occupied by a single cell (Fig. 2b, cross-section: middle region, pink cell; dorsal region, blue cell). This uneven cell distribution persists and is consistently observed from 3 to 4 dpf (Supplementary Fig. 3a & b and Supplementary Movie 2). This observation supports our strain analysis, which demonstrated an inverse trend in ES at both radial and vertical axes (Fig. 1e) as a consequence of cell rearrangement. Timelapse imaging confirmed this and showed that this inverse trend is driven by cell rearrangements and uneven cell occupation of the circumferential surface.

In contrast to aISVs, all vISVs examined remained multicellular between 2 and 3 dpf, with three or more cells maintaining cell-cell junctions to form characteristic “Y-shaped” junctions (Fig. 2c). The phenomenon of vessels being wrapped by a single cell is rarely observed in vISVs. Occasionally, vISVs exhibit X-shaped junctions, but these do not elongate into self-seam junctions. Instead, ECs in vISVs actively divide, as indicated by the accumulation of F-actin in the rounded mitotic cell (Fig. 2c, 62 hpf, black arrows, & Supplementary Movie 3). To accommodate the increased cell number, ECs significantly decrease in size, followed by the dorsal migration of one daughter cell that contributes to the DLAV (Fig. 2c). Such dorsal cell rearrangements and cell division continue to be observed between 3 and 4 dpf, though no further contributions to the DLAV were noted (Supplementary Fig. 3b & Supplementary Movie 4).

These data indicate that ECs dynamically control their size, shape and position to remodel blood vessels, leading to vessel constriction and elongation and redistribution of EC number. While strain analyses from static time points (across days) suggest that aECs exhibit less cell deformation as compared to vECs, continuous time-lapse imaging (across hours) revealed that aECs undergo highly dynamic shape changes. To maintain the integrity of the vessel lumen while adjusting their positions, aECs with low cell numbers employ a mechanism where a single cell wraps around the entire vessel, facilitating the rapid movement of other cells along the vessel. Subsequently, these aECs reduce in size and migrate vertically to reduce cell occupancy at the radial axis, resulting in uneven cell distribution on the circumferential surface, collectively contributing to a reduction in vessel size. In contrast, vISVs are primarily multicellular tubes with a greater number of cells (at 3 dpf, vISVs consist of an average of 9 ECs while aISVs consist of 5), where vECs significantly decrease in size to accommodate the increased cell count within the vessel.

### Transitions in cortical actin organization during vessel constriction

To investigate the role of the actin cytoskeleton remodelling in the process of vessel constriction, we monitored the endothelial actin reporter line with high-resolution microscopy. We identified distinct cortical actin structures, which we classified into three types: circumferential (C), mesh (M), and longitudinal (L) actin organizations (AOs, Fig. 3a). At 2 dpf, we observed a mixture of all three AOs. From 3 dpf onward, there was a reduction in circumferential and mesh AOs, accompanied by an increase in longitudinal AO. By 4 dpf, longitudinal AO becomes the dominant structure. This AO transition is conserved in both aISVs and vISVs, although at 2 dpf, aISVs contain more circumferential AO, whereas vISVs exhibit a greater presence of mesh AO by 4 dpf (Fig. 3b).

**Figure 3.**
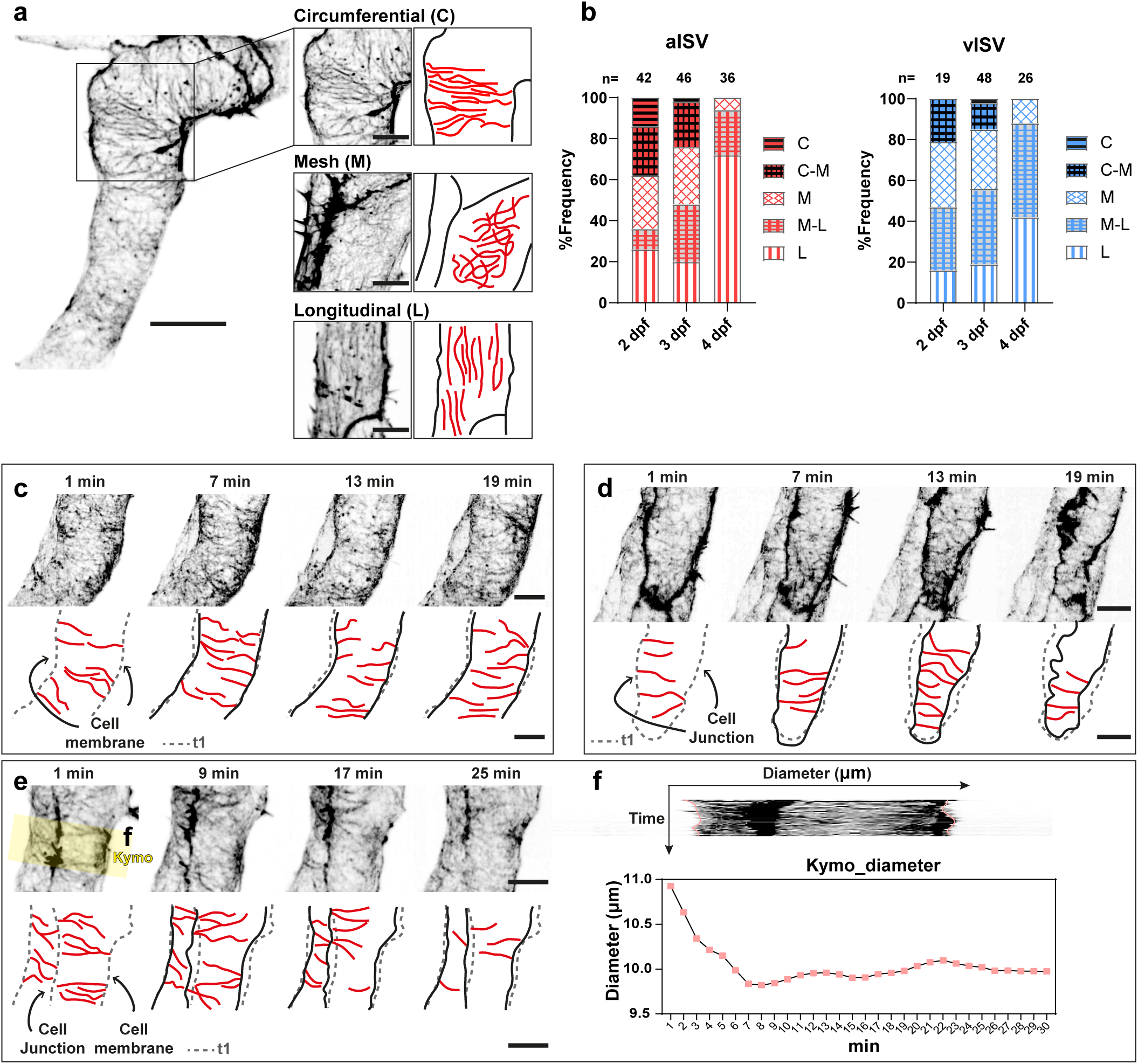
Distinct actin organization transition and the correlation between circumferential actin and vessel constriction. **a** Representative images of circumferential, mesh, and longitudinal actin organizations. Scale bar, 10 μm. **b** Percentage of each actin organization type from 2 dpf to 4dpf in aISVs and vISVs. Total number of ISVs (from 5 independent experiments) is indicated on the top of the bar. **c-e** Still frames from time-lapse movies (Supplementary Movies 5-7) illustrating circumferential actin anchoring to the cell membrane (**c**), connecting to cell-cell junctions (**d**), or linking both the membrane and cell junctions (**e**). In the schematics, circumferential actin is highlighted in red. Black lines trace the cell boundary at the current time point, and grey serrated lines represent the outline of the cell membrane or cell-cell junctions at the first time frame for comparison. Scale bar, 5 μm. **f** Kymograph from the yellow ROI in **e**, and measurements of diameter over time. ISV intersegmental vessel; aISV arterial ISV; vISV venous ISV.

We next investigated the spatiotemporal relationship between actin organization and cell deformation. To achieve this, we performed fast time-lapse imaging (1-minute intervals) of F-actin to track the dynamics of both cortical and junctional actin. Our findings revealed that circumferential actin can anchor to the cell membrane (Fig. 3c & Supplementary Movie 5); connect to two cell-cell junctions (Fig. 3d & Supplementary Movie 6); or connect to a cell-cell junction and the membrane (Fig. 3e & Supplementary Movie 7). Notably, the connection of actin bundles with cell junctions and/or plasma membranes coincides with inward cell deformation (Fig. 3c – e, illustrations). We additionally measured vessel diameter and found that when there are long and persistent circumferential actin, the vessel diameter decreases (Fig. 3f). These findings suggest that circumferential actin plays a key role in driving cell contraction and vessel constriction.

### Circumferential actin, together with myosin II, transmits the strongest tensile force

Our observations suggest that circumferential actin may mediate contractile forces, potentially generated by non-muscle myosin II (hereafter, referred to as myosin II), to drive vessel constriction. To examine this hypothesis, we used the transgenic lines *Tg(fli1:GAL4FF)^ubs^*^3^ *; Tg(UAS:EGFP-UCHD)^ubs^*^18^ and *Tg(CxUAS:mylSb-mCherry)^rk^*^32^ to visualize actin and myosin II, respectively. Time-lapse imaging demonstrated the co-localization of actin and myosin II, with myosin II largely presenting as punctate structures along actin bundles throughout the 20-minute imaging window. Notably, one instance of linear myosin II alignment along circumferential actin bundles is observed (Fig. 4a & Supplementary Movie 8). Further, spatial co-localization of actin and myosin II at the 8-minute timepoint was confirmed by 3D rendering (Fig. 4b & Supplementary Movie 9). Kymograph of regions exhibiting linear *myl9b-mCherry* expression indicated that increased actin and myosin II intensity is associated with reduced vessel diameter (Fig. 4c). This pattern of linear co-localization appeared consistently across all time-lapse recordings (n = 20/20) where circumferential actin is present.

**Figure 4.**
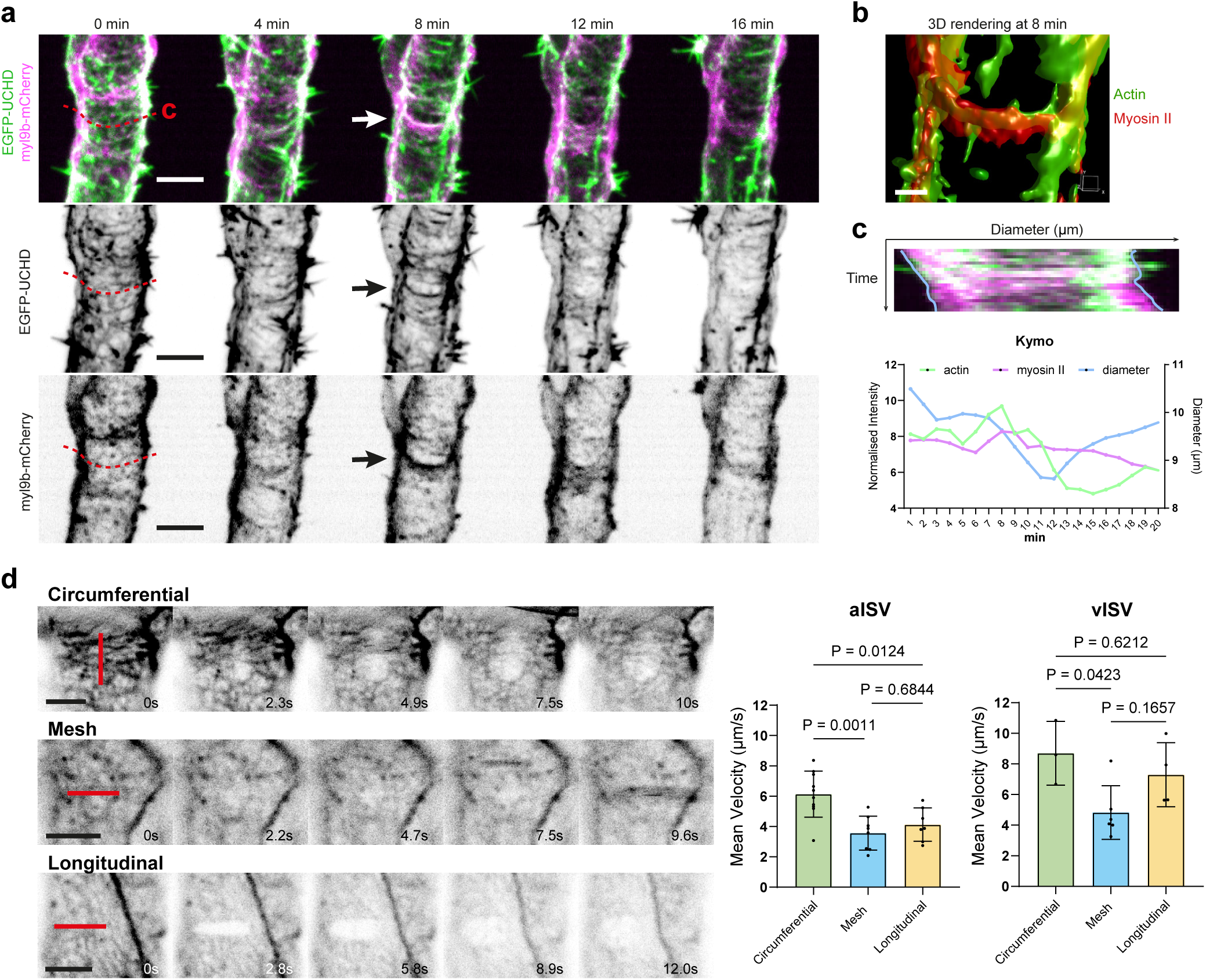
Tensile force transmission by linear co-localization of circumferential actin and myosin II in vessel constriction. **a** Still images (Supplementary Movie 8) of an aISV at 2dpf showing actin (EGFP-UCHD) and non-muscle myosin II (myl9b-mCherry) from *Tg(fli1:GAL4FF)^ubs^*^3^ *; Tg(UAS:EGFP-UCHD)^ubs^*^18^ and *Tg(CxUAS:mylSb-mCherry)^rk^*^32^. Linear colocalization of circumferential actin and myosin II is observed at 8 min. Similar observations were seen in 20 movies. **b** 3D rendering (Supplementary Movie 9) of actomyosin colocalization at 8 min showed in **a**. **c** Kymograph of region indicated in **a** and measurements of normalised average fluorescent intensity of UCHD and myl9b and vessel diameter. **d** Laser ablation (red line) on circumferential, mesh, and longitudinal actin organisation (Supplementary Movie 10-12). Mean velocity from the first 10s right after laser ablation is measured (aISV n=10/8/7 and vISV n=3/6/4 in C/M/L). Statistical significance was assessed by ordinary one-way ANOVA with Turkey’s multiple comparisons test. Mean values are indicated. ISV intersegmental vessel; aISV arterial ISV; vISV venous ISV.

To directly assess whether the three actin organizations bear tensile forces, we conducted laser ablation experiments using a multiphoton microscope. Ablation of circumferential actin resulted in a rapid recoil of the actin bundles, creating a gap in the cortical network (Fig. 4d). The mean recoil velocity is highest for circumferential actin, followed by longitudinal actin, with mesh actin organization exhibiting the least recoil velocity (Fig. 4d). These results indicate that circumferential actin transmits the strongest tensile forces.

Collectively, our findings demonstrate that the actin bundles, especially circumferential actin, serve as a scaffold for myosin II contractile activity that generates forces that pull on cell junctions and membranes. This contractile activity drives cell deformation, thereby facilitating cell contraction and cell rearrangement and contributing to vessel constriction.

### Imbalance in the proportion of circumferential and mesh actin compromises vessel constriction

Our findings indicate that circumferential actin plays a crucial role in generating contractile forces necessary for vessel constriction. To further support this notion, we modified the relative proportion of actin organization by experimentally increasing the prevalence of mesh actin through the overexpression of WASp, a key regulator of actin nucleation that facilitates the formation of branched actin networks^10,23^. A plasmid encoding *6xUAS:wasb-mCherry* was selectively expressed in ECs in *Tg(fli1:GAL4FF)^ubs^*^3^*; Tg(UAS:EGFP-UCHD)^ubs^*^18^ embryos (Fig. 5a & d). As expected, the overexpression (OE) of *wasb* specifically in ECs resulted in an increase in mesh actin organization, and therefore a reduction in circumferential actin organization in both aISVs and vISVs at 2, 3 and 4 dpf (Fig. 5a – c) when compared to control (*wasb-mCherry* negative) ECs. This alteration is associated with a significant increase in vessel diameter in both aISVs and vISVs at 2, 3 and 4 dpf (Fig. 5d – f) and reduced vessel constriction (17% reduction of diameter in control and 14% in OE-vessels) during this period when compared to control, thereby supporting our hypothesis that circumferential actin is essential for vessel constriction.

**Figure 5.**
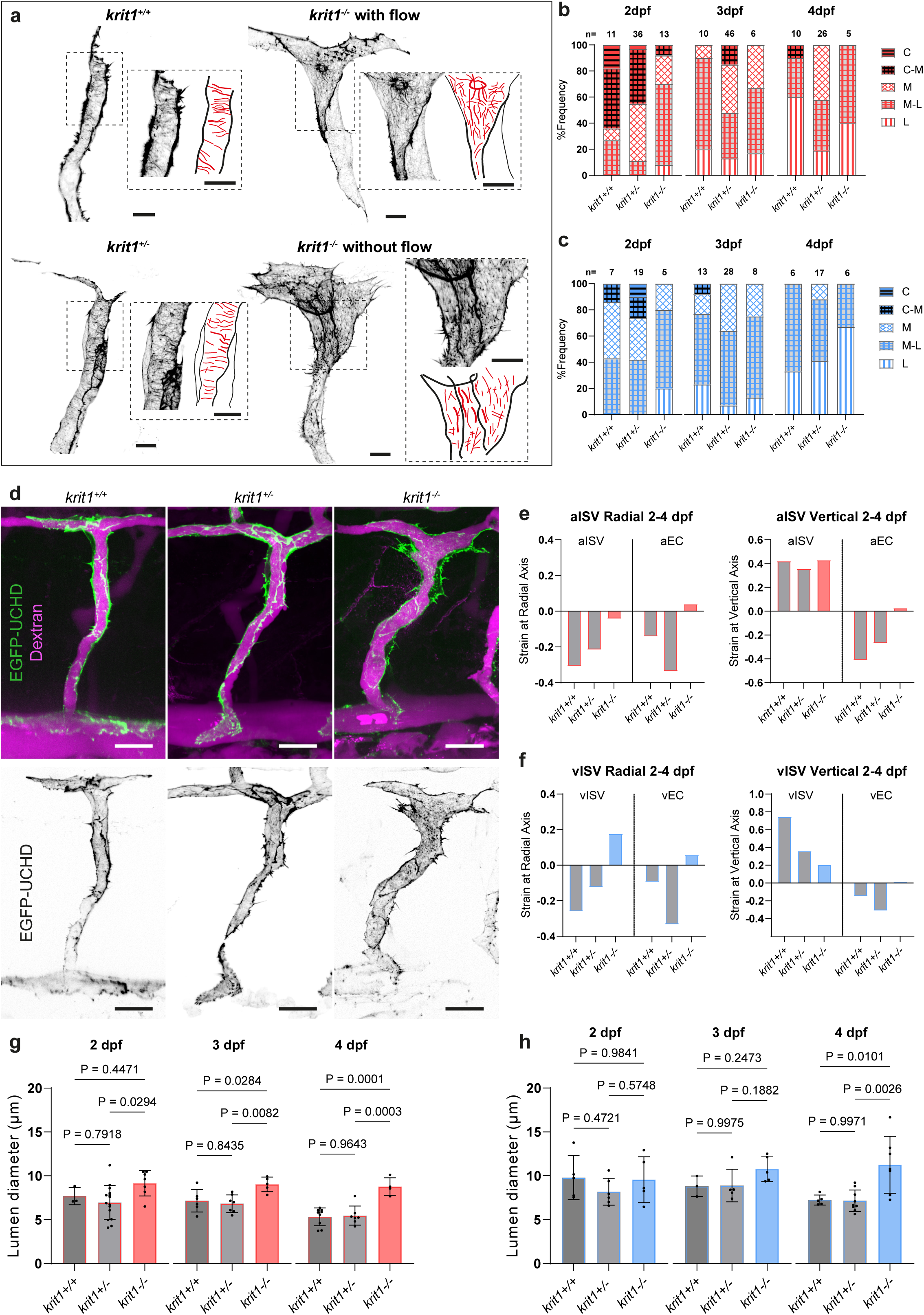
Manipulation of actomyosin disrupts vessel constriction. **a**, **d** Representative images of ISVs of 2 dpf (*Tg(fli:Gal4ff^ubs^*^3^*;UAS:EGFP-UCHD^ubs^* embryos expressing constructs encoding *CxUAS:wasb-mCherry*. Actin organizations are highlighted in red in the schematics. Scale bar, 5 μm (**a**) and 20 μm (**d**). **b**, **c** Percentage of actin organisation between control (wasb-mCherry negative) and wasb-mCherry overexpressing cells from 2 dpf to 4 dpf in aISVs and vISVs. Total number of ISVs is indicated on the top of each bar. **e**, **f** Quantification of vessel diameter between control (wasb-mCherry negative) and wasb-mCherry overexpressing cells from 2 dpf to 4 dpf in aISVs and vISVs. Each data point represents one ISV. **g** Representative images of ISVs of 2 dpf *Tg(fli1:myr-mCherry)^ncv^*^1^ embryos expressing constructs encoding dominant negative myosin light chain protein *CxUAS:mylSbA2A3-eGFP*. Scale bar, 20 μm. **h**, **i** Quantification of vessel diameter between control (*mylSbA2A3-eGFP* negative) and *mylSbA2A3-eGFP* overexpressing cells from 2 dpf to 4 dpf in aISVs and vISVs. Each data point represents one ISV. Statistical significance was determined by two-tailed unpaired *t-test*. **j** Representative images of ISVs of 2 dpf wildtype embryos expressing constructs encoding dominant negative myosin light chain protein *CxUAS:mylSbA2A3-eGFP* in single cells. Microangiography is performed by injecting dextran rhodamine. Scale bar, 20 μm. Statistical significance was determined by two-tailed unpaired *t-test*. **k** Strain analysis at radial and vertical axis between 2-4 dpf in aISVs between control (radial: aISV, n=39/37, aEC n=16/32 at 2/4 dpf; vertical: aISV n=24/30, aEC n=16/32 at 2/4 dpf) and myl9bA2A3A-OE (radial: aISV, n=32/41, aEC n=24/18 at 2/4 dpf; vertical: aISV n=23/28, aEC n=24/18 at 2/4 dpf). **l** Strain analysis at radial and vertical axis between 2-4 dpf in vISVs between control (radial: vISV, n=29/33, vEC n=13/21 at 2/4 dpf; vertical vISV n=17/26, vEC n=13/21 at 2/4 dpf) and myl9bA2A3-OE (radial: vISV, n=23/36, vEC n=12/27 at 2/4 dpf; vertical vISV n=23/33, vEC n=12/27 at 2/4 dpf). ISV intersegmental vessel; aISV arterial ISV; vISV venous ISV. Scale bars, 20 µm. Source data are provided as a Source data file.

### Inhibition of myosin II activity leads to disrupted cell deformation and increased vessel diameter

To investigate the impact of reduced myosin II activity on EC deformation and vessel diameter, we overexpressed a dominant-negative myosin light chain protein (Myl9bA2A3) tagged with eGFP in *Tg(fli1:myr-mCherry)^ncv^*^1^ in a mosaic manner (Fig. 5g & j). aISVs and vISVs with *myl9bA2A3* overexpression (*myl9bA2A3-OE*) exhibit increased diameters from 2 to 4 dpf (Fig. 5H & I). To further elucidate the effect of *myl9bA2A3-OE* on cell deformation, we conducted single-cell shape and strain analyses (Fig. 5j). The cell areas of *myl9bA2A3-OE* cells increase at 4 dpf in both aISVs and vISVs (supplementary Fig.4a Cb). Moreover, aISVs composed of *myl9bA2A3-OE* cells exhibit a smaller negative radial tissue strain when compared to control aISVs, suggesting that while myl9bA2A3-OE vessels constrict over time, the degree of constriction between 2 and 4 dpf is reduced (Fig. 5k). Notably, cell strain in myl9bA2A3-OE aISVs is positive along both the radial and vertical axes while that of control aECs is negative, indicating an expansion in cell dimensions (Fig. 5k). On the other hand, vISVs composed of *myl9bA2A3-OE* cells exhibit similar strain along the radial and vertical axes compared to control. However, while cell strain in control vECs become increasingly negative at both radial and vertical axes over time, *myl9bA2A3-OE* cells do not undergo significant reduction in cell strain especially in the radial axis, implying impaired contractility due to disrupted myosin II activity (Fig. 5l). These findings thus highlight the critical role of the actomyosin network in governing cell deformation and vessel constriction.

### Loss of circumferential actin in *krit1^-/-^* endothelial cells results in increased cell size and vessel diameter

Our results above demonstrate a role of actomyosin-driven EC deformation in blood vessel remodelling to establish appropriately sized vessels. To further explore this, we investigated whether compromised actomyosin activity might underlie the vascular malformations observed in CCM. Prior research has demonstrated that the loss of *ccm1*/*krit1* or *ccm2* in zebrafish causes major blood vessels to dilate and results in unregulated cell size and shape^17,18^. To investigate whether altered formation of actin cytoskeleton contributes to CCM-related vascular malformations, we examined *krit1* mutant zebrafish. However, assessing actin-dependent deformations in *krit1* mutant zebrafish is challenging due to cardiac defects and subsequent disrupted blood flow^18^, which significantly impact actin dynamics and cell shape^24^. Consequently, we performed cell transplantation experiments to investigate the autonomous role of Krit1 in regulating actin organization and EC shape. By in-crossing *krit1^+/t2C^*^458^*; Tg(fli1:GAL4FF)^ubs^*^3^*; Tg(UAS:EGFP-UCHD)^ubs^*^18^ lines and using the resulting embryos as donors for transplantation into wild-type recipients, we analysed actin organization, cell deformation, and vessel lumen diameter within endothelial cell grafts.

The results showed that wild-type and *krit1^+/t2C^*^458^ (hereafter referred to as *krit1^+/-^*) ECs exhibit both circumferential and mesh actin organizations from 2 dpf (Fig. 6a-c). In contrast, *krit1^-/-^* ECs completely lack circumferential actin bundles, with longitudinal actin predominating from 2 to 4 dpf in both aISVs and vISVs (Fig. 6a-c). In some grafts with disrupted blood flow, *krit1^-/-^* ECs become markedly flattened, enlarged, and misaligned, displaying dominant longitudinal actin and straightened cell-cell junctions (Fig. 6a). The absence of circumferential actin in *krit1^-/-^* ECs correlates with defects in cell deformation and vessel constriction (Fig. 6d). Rather than undergoing negative cell strain as observed in wild-type and *krit1^+/-^* aECs and vECs, *krit1^-/-^* ECs exhibit slightly positive cell strain along the radial axis while remaining unchanged along the vertical axis (Fig. 6e, f). These findings suggest that ECs in CCMs fail to contract. Indeed, the area of *krit1^-/-^* ECs is increased (supplementary Fig. 5a & b). Consequently, *krit1*^-/-^ EC-containing aISVs and vISVs have significantly enlarged lumens at 2, 3 and 4 dpf when compared to wild-type and *krit1^+/-^* EC-containing vessels (Fig. 6g, h). Importantly, the progressive constriction of vessel diameter observed in wild-type and heterozygous vessels was lost in *krit1^-/-^*. Instead, vessel strain is less negative in aISVs and more positive in vISVs at the radial axis (Fig. 6e, f). These findings collectively indicate that Krit1 regulates cell and vessel size autonomously and that the lack of circumferential actin formation in *krit1^-/-^* ECs impairs cell deformation and prevents vessel constriction, thereby manifesting in vessel malformation.

**Figure 6.**
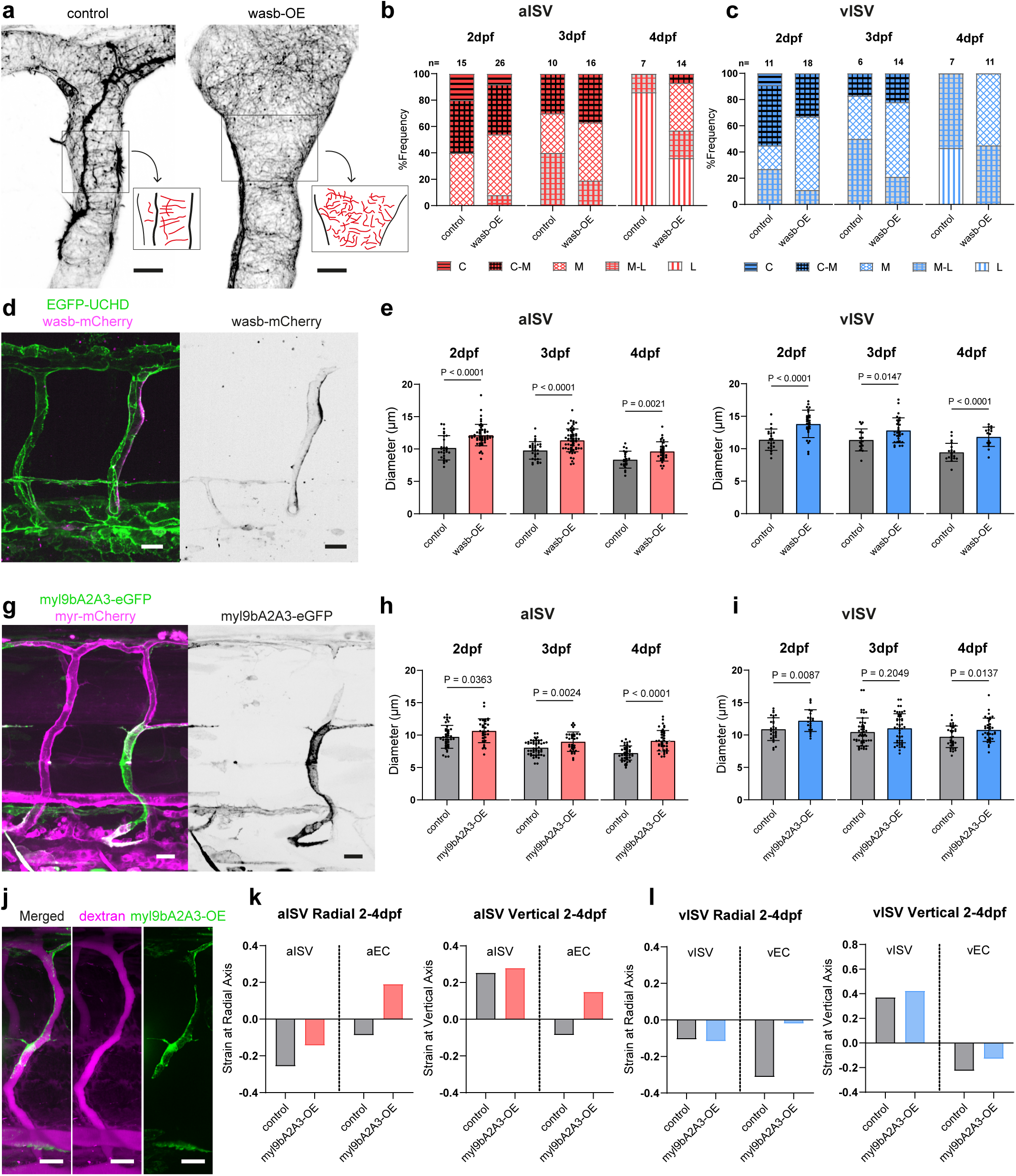
Loss of circumferential actin in *krit1^-/-^* endothelial cells disrupts cell deformation and vessel constriction. **a** Representative images of transplanted ISVs of 2 dpf *krit1^+/t2C^*^458^; (*Tg(fli:Gal4ff^ubs^*^3^*;UAS:EGFP-UCHD^ubs^* into wildtype embryos, showing different actin organisations (highlighted in red in schematics). Scale bar, 10 μm. **b**, **c** Percentage of actin organisation between *krit1^+/+^*, *krit1^+/-^* and *krit1^-/-^* cells from 2 dpf to 4 dpf in aISVs and vISVs. Total number of ISVs is indicated on the top of each bar. **d** Microangiography in transplanted wild type and *krit1* mutant ISVs. Scale bar, 20 μm. **e** Strain analysis at radial and vertical axis between 2-4 dpf in aISVs between *krit1^+/+^* (aISV n=3/9, aEC n=4/13 at 2/4 dpf), *krit1^+/-^* (aISV n=15/7, aEC n=17/14 at 2/4dpf) and *krit1^-/-^* (aISV n=7/5, aEC n=15/5 at 2/4 dpf). **f** Strain analysis at radial and vertical axis between 2-4 dpf in vISVs between *krit1^+/+^* (vISV n=5/5, vEC n=5/8 at 2/4 dpf), *krit1^+/-^* (vISV n=6/9, vEC n=10/20 at 2/4 dpf) and *krit1^-/-^* (vISV n=5/7, vEC n=7/13 at 2/4 dpf). **h**, **i** Quantification of lumen (indicated by dextran) diameter between *krit1^+/+^*, *krit1^+/-^*, *krit1^-/-^* transplanted cells from 2 dpf to 4 dpf in aISVs and vISVs. Each data point represents one ISV. Statistical significance was assessed by ordinary one-way ANOVA with Turkey’s multiple comparisons test. Mean values are indicated.

## Discussion

The tight control of blood vessel size is crucial for efficient blood flow distribution and perfusion of tissues. During embryonic development, vessel size is determined during the period of vascular remodelling. By performing high-resolution, multiscale study of actin cytoskeleton organization, EC shape and vessel diameter in developing zebrafish embryos at different stages of development, we demonstrate that vessel remodelling is a dynamic process requiring the coordination of EC shape changes, cell rearrangements and cell division, collectively driving both vessel constriction and elongation. Our detailed analyses of cortical actin organization and dynamics and myosin II function uncovered a key function of circumferential actomyosin bundles in deforming ECs to constrict blood vessels. When this contractile machinery fails to form, as demonstrated in ECs with reduced myosin II activity and Krit1 function, ECs do not deform, leading to dilated vessels that are characteristic of CCMs.

Vasoconstriction is thought to mainly result from the activity of mural cells, which can be regulated by ECs through paracrine signalling to control vascular tone. Mural cells, including vascular smooth muscle cells (vSMCs) and pericytes, wrap around ECs, contribute to vessel diameter regulation through their contractile activity and/or their role in producing and stabilising vascular basement membrane^25–28^. However, in zebrafish, pericyte ensheathment of ISVs occurs later, after 60 hpf, when remodelling has already started, and only cover 50% vISVs by 120 hpf^29^, suggesting that they may not be essential for initiating vessel constriction and underscores the importance of endothelial regulation of vessel diameter during vascular remodelling.

Our study highlights that ECs can directly regulate vessel constriction via intrinsic mechanisms, specifically through shape changes and deformation mediated by circumferential actin. However, we also found that venous ECs shrink more than arterial ECs, despite the latter exhibiting more circumferential actin at 2 dpf, suggesting the underlying mechanisms may differ between vessel types. This difference is likely due to variations in EC numbers. Given the physical constraints and the production of smaller daughter cells through division^30^, venous ECs, with their higher division rates, may require less circumferential actin-mediated deformation (Fig. 7). In contrast, arterial ISVs primarily acquire ECs through migration from the DLAV, a process that requires significant circumferential actin-mediated deformation to accommodate new cells and facilitate vessel elongation and narrowing (Fig. 7).

**Figure 7.**
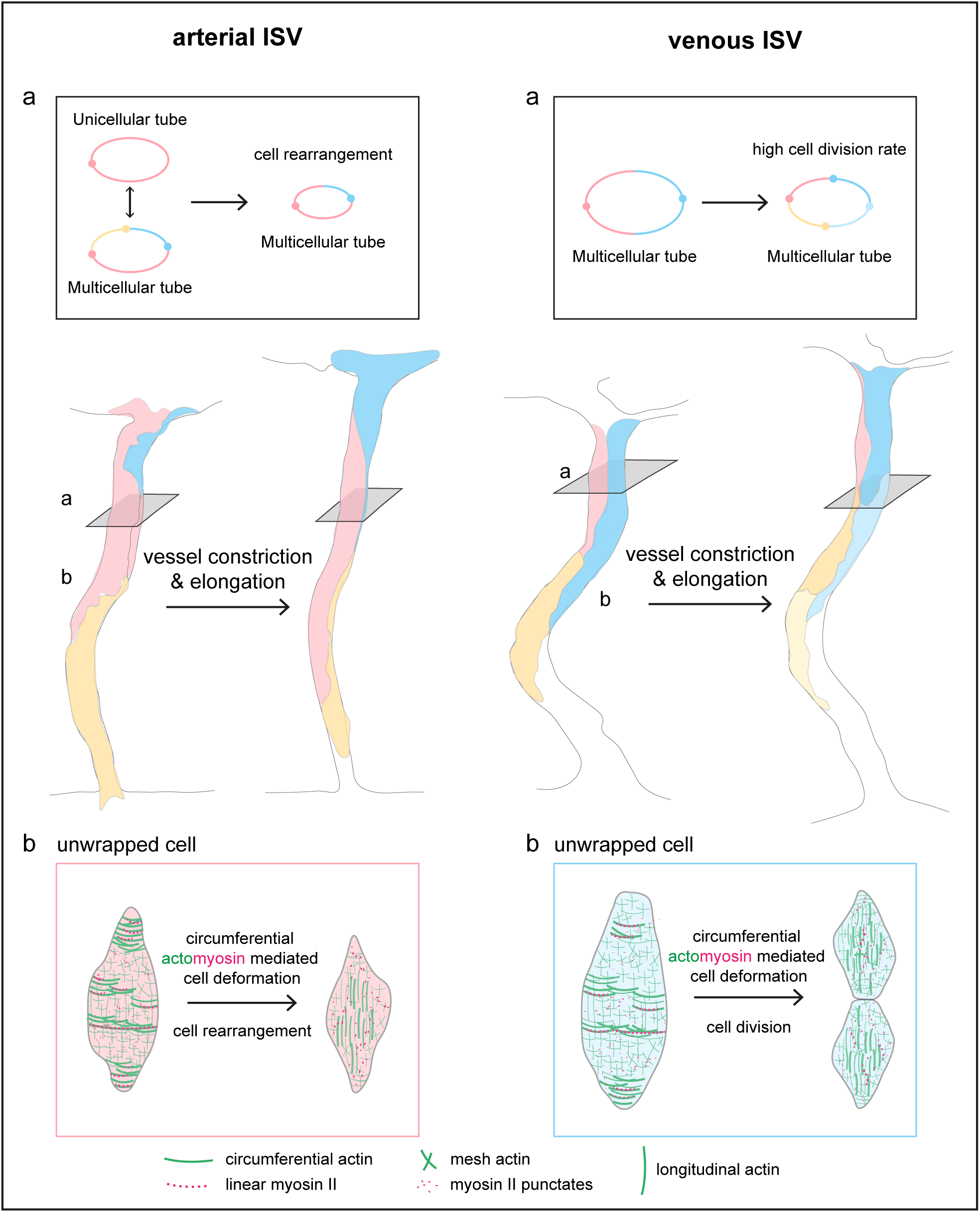
Graphic model illustrating the mechanisms of arterial and venous ISV remodelling during development. Arterial ISVs and venous ISVs undergo extensive remodelling during development, including elongation and constriction. In the aISVs (left panel, a), endothelial cells undergo dynamic cell rearrangements, characterized by the transient formation of self-seam junctions. This process enables the vessel to transition between unicellular and multicellular tube configurations, highlighting a critical mechanism by which changes in EC rearrangement regulate vessel diameter. On the other hand, vISVs (right panel, a) experience extensive EC divisions and to accommodate an increased number of cells, ECs undergo a marked reduction in size (right panel, b). EC deformations and rearrangements contribute to a negative relationship between cell number and vessel diameter. These deformations are driven by circumferential actomyosin contraction, which facilitates cell shape changes and migration to ensure proper vessel remodelling (b).

By establishing high-resolution timelapse imaging of actin cytoskeleton at different stages of development, we not only reveal the presence of different actin organizations in the EC cortex but also show that the actin network remodels over time. This represents, to our knowledge, the first *in vivo* high-resolution, temporal analysis of endothelial actin organization, at a level of detail typically only achievable in *in vitro* studies. How actin organization and remodelling in ECs are regulated during vessel remodelling remains to be determined. It is likely that molecular regulators of actin cytoskeleton play a key role in the organization of actin bundles. It has been demonstrated in *Drosophila* trachea that the formin actin nucleator protein, DAAM, helps align actin bundles along the circumferential direction^31,32^. In this study, we show that the ectopic expression of wasb in ECs reduces the proportion of circumferential and longitudinal actin in favour of more mesh actin. Actin organization in ECs can also be regulated by external mechanical stresses. In *in vitro* 3D vascular tubes, the elevation in luminal pressure (from 150 Pa to 650 Pa) reorganizes actin cytoskeleton from a longitudinal to circumferential orientation^21^. ECs are also responsive to fluid shear stress. It is well established from 2D *in vitro* experiments that fluid shear stress induces the alignment of actin bundles in the direction of flow that increases under strong laminar flow^33^. In zebrafish ISVs, circumferential actin is most abundant at 2 dpf and diminishes thereafter. Our previous study shows that blood flow velocity and wall shear stress increase from 2 to 3 dpf in aISVs, after which there is a continuous decline until 6 dpf^34^. Hence, the relationship between luminal forces – pressure and wall shear stress – on endothelial actin cytoskeleton *in vivo* is more ambiguous. The overall actin organization *in vivo* is most likely determined by the combined effects of stretch, wall shear stress as well as the state that the ECs are in (for example, whether the cell is migratory, proliferative etc.). Furthermore, the effects of basement membrane deposition and pericyte ensheathment on EC actin organization remains to be explored.

In addition to the *de novo* actin organization observed *in vivo*, we identified the formation of a temporary self-seam junction during vessel remodelling. This phenomenon, where a single cell forms self-contacts, resembles Type I vessel anastomosis, wherein a cell encircling a unicellular tube splits on one side to enable neighbouring cell contacts. However, unlike self-splitting, the self-seam junction persists for hours without resolving. Interestingly, this mechanism aligns more closely with “self-fusion,” as described by Lenard et al. (2013), where a cell wraps around a vessel, forms temporary contacts, and extends its membrane to create a seamless tube. In contrast, the self-seam junction remains stable, representing an intermediate state between self-splitting and self-fusion. This self-contact mechanism appears to facilitate efficient transitions from a unicellular tube back to a more stable multicellular tube configuration. Similar remodelling of tri-cellular junctions, driven by actomyosin and promoting cell rearrangement, has been observed in epithelial morphogenesis in *Drosophila*^35^. This transient self-contact mechanism likely serves a critical function in maintaining vessel integrity amidst the dynamic and mechanically demanding processes of vascular remodelling.

While previous studies on CCMs have uncovered key signalling pathways that are dysregulated in the disease^4^, it is still unclear how altered signalling causes the pathological vessel morphology. By taking advantage of the tractability of the zebrafish to perform high-resolution imaging, single cell analysis and time-course experiments, we have now gained further insights into the cellular behaviour of Krit1-deficient ECs during vessel remodelling. We demonstrate for the first time that ECs lacking Krit1 exhibit a reduced capacity for cortical actin remodelling and cell deformation. The inability of Krit1-deficient ECs to remodel cortical actin effectively emphasizes the critical role of actin-driven cell shape changes in mediating vessel constriction. Mechanistically, Krit1/CCM1 and CCM2 form protein complexes that stabilize the connections between junctional proteins, such as VE-cadherin, and the actin cytoskeleton through interactions with Rap1 and β-catenin, as well as integrins via ICAP-1^36–38^. The loss of Krit1 likely disrupts these stabilizing interactions, leading to aberrant actin organization and impaired actomyosin-mediated contractility. While studies show that Krit1-deficient ECs display enhanced actomyosin-mediated contractility and stress fibre formation as a result of upregulated ROCK signalling^38–40^, Yin et al demonstrated in a recent study that there is reduced myosin II-mediated contractility along endothelial junctions in *krit1* zebrafish mutants^41^. Together with our observed defects in actin organization and remodelling in *krit1* zebrafish mutants, the latter finding supports a critical function of actomyosin-mediated contractility in deforming ECs to constrict blood vessels. Whether the longitudinal actin structures in Krit1-deficient ECs we observed are stress fibres remains unclear and needs further examination. Interestingly, wild-type ECs predominantly exhibit circumferential actin, whereas Krit1-deficient ECs undergo a notable shift toward longitudinal actin organization. This transition from circumferential to longitudinal actin may stem from increased sensitivity of Krit1-deficient ECs to shear stress, which preferentially aligns actin fibres in the direction of blood flow. Restoration of blood flow in *krit1* mutant zebrafish protects large arterial vessels from vascular anomalies but fails to prevent abnormalities in smaller brain vessels^18^. Consistently, our study shows that both arterial and venous ISVs exhibit abnormal EC size and shape despite maintained blood flow. ECs in regions without flow show even greater enlargement, suggesting that low flow exacerbates *krit1* dysfunction. These findings indicate that slow flow may serve as a key trigger for *ccm1* dysfunction. Additional factors likely contribute to the differential vulnerability of large and small vessels including extracellular matrix remodelling^39^ and pericyte regulation^42^.

In conclusion, our investigations have uncovered the mechanics how of blood vessel diameter is controlled through coordinated EC deformations and cell rearrangements. Specifically, we have identified an essential role of circumferential actomyosin bundles in deforming ECs in the radial axis to constrict vessels during vascular remodelling. Such actomyosin-induced cell shape transition is disrupted in a zebrafish model of CCM, providing a cellular mechanistic understanding of how vascular malformations arise and implicates impaired EC mechanics as an underlying factor in the aetiology of vascular anomalies in which vessels are abnormally dilated.

## Methods

### Zebrafish husbandry and manipulation

Zebrafish (*Danio rerio*) were maintained and raised following standard protocols. The transgenic lines used in this study included *Tg(fli1:MYR-EGFP)^ncv^*^2^^43^, *Tg(fli1:h2bc1-mCherry)^ncv^*^31^^44^*, Tg(fli1:myr-mCherry)^ncv^*^1^^45^, *Tg(fli1:GAL4FF)^ubs^*^3^^46^, *Tg(UAS:EGFP-UCHD)^ubs^*^18^^47^ *, Tg(CxUAS:mylSb-mCherry)^rk^*^32^ (this study), while the mutant line used was *krit1^t2C^*^458^^17^. All animal experiments were approved by the Institutional Animal Care and Use Committee at RIKEN Kobe Branch (IACUC). Fish embryos were collected within 30 minutes of divider removal and were allowed to develop at 28.5°C to the appropriate stage. For genotyping of the *krit1^t2C^*^458^ mutant, genomic DNA was extracted using the HotSHOT method, and PCR amplification was performed with specific primers (Forward 5’-CCACAAGCGTAACGTAAATG; Reverse 5’-ATCTATGGACGCAATGCAG). The resulting PCR products were purified and sequenced using the forward primer.

### Plasmid production and injection

Plasmids used in this study were constructed using either the NEBuilder® HiFi DNA Assembly or In-Fusion® HD Cloning kits, following the respective manufacturer protocols. The *wasb-mCherry* vector^10^ (a gift from Holger Gerhardt, Max Delbrück Center for Molecular Medicine, Germany) was amplified by PCR and incorporated into Ac/Ds transposon-based constructs. Similarly, the *myl9bA2A3-eGFP* vector^48^ (a gift from Anne Schmidt, Institut Pasteur, France) was amplified by PCR and assembled into Tol2 transposon-based constructs. Assembly reactions employed the appropriate backbone vectors and primers as specified by the cloning protocols. For details on vectors and primers, refer to the supplementary tables. For plasmid overexpression experiments, embryos were injected at the one-cell stage with 2 nl of a mixture containing 50-100 ng of plasmid DNA, 100-200 ng Tol2 or Ac transposase mRNA, phenol red for tracking, and Milli-Q water to a final volume.

### Imaging

Embryos were immobilized in 0.8% low-melt agarose (Bio-Rad) within E3 medium supplemented with 0.16 mg/mL Tricaine and 0.003% phenylthiourea to reduce movement and pigmentation. Confocal imaging was performed using an inverted Olympus IX83 microscope, equipped with a Yokogawa CSU-W1 spinning disk confocal unit. Imaging was carried out with a 60x/1.2 NA water immersion objective (Olympus UPLSAPO) and a Zyla 4.2 CMOS camera (Andor). Image acquisition was controlled using Andor iQ3 software, and z-stack images were captured for 3D reconstructions. Actin organization was imaged using two advanced microscopy setups. The first system was an Andor/Olympus Dragonfly 200 inverted microscope equipped with a 60x water immersion objective lens (Olympus UPLSAPO, NA 1.2) and a Zyla 4.2 PLUS sCMOS camera (2.0x magnification). Imaging utilized a 40 µm pinhole, and z-stacks were acquired at Nyquist intervals. The second system was a customized spinning disk superresolution microscope (SDSRM)^49^, implemented based on the commercial IXplore IX83 SpinSR system (Evident), equipped with a 60x silicon immersion objective lens (Evident UPLSAPO 60XS, NA 1.3). z-stack interval was 0.15um.

### Microangiography

Microangiography was performed for single cell analysis. Embryos at 2, 3, and 4 dpf were injected with 1–2 nL of dextran tetramethyl rhodamine (molecular weight 2000 kDa, Invitrogen) at a concentration of 10 mg/mL. The dextran solution was introduced through the duct of Cuvier of the zebrafish. Imaging was performed immediately following the injection. Confocal z-stacks were acquired using an Olympus UPLSAPO 60x/1.2 NA water immersion objective.

### Cell transplantation

Donor cells from embryos derived from *krit1^+/t2C^*^458^ in-crossed zebrafish in *Tg(fli1:GAL4FF)^ubs^*^3^*; Tg(UAS:EGFP-UCHD)^ubs^*^18^ background were collected from random locations in the blastoderm. These cells were transplanted into the lateral marginal zone of the blastoderm in wild-type recipient embryos at 4.5–6 hpf. The procedure was carried out using a CellTram® 4r Oil microinjector (Eppendorf) with a borosilicate glass capillary GC100-15 (Harvard Apparatus Ltd). The capillary tips were shaped into a smooth “spoon” using a horizontal micropipette puller P-87 (Sutter Instrument Co.) and a MF-900 microforge (Narishige) for precision. During transplantation, embryos were maintained in agarose-coated Petri dishes containing 0.5x E2 buffer composed of 7.5 mM NaCl, 0.25 mM KCl, 0.5 mM MgSO_4_, 75 µM KH_2_PO_4_, 0.5 mM CaCl_2_, 0.35 mM NaHCO_3_, and 50 U/mL penicillin-streptomycin. Recipient embryos were grown to 2, 3, and 4 dpf, and those exhibiting EGFP-positive ISVs were selected for microangiography and imaging. Genotyping was performed on the corresponding donor embryos. A total of 17 independent transplantation experiments was conducted.

### Image processing

Raw images of actin organization captured using the Dragonfly 200 microscopy were deconvoluted with ™Huygens Spinning Disk Deconvolution, following the standard protocol established in the unit. The deconvoluted images were subsequently processed using Fiji software (NIH). To separate the front and back sides of the vessel, the ISV was first straightened using Fiji’s *Straighten* tool and then divided with the *Split* tool. Z-stacks for each half were projected into single images using *maximum intensity projection.* To enhance consistency and reproducibility, a custom macro was developed in Fiji to semi-automate the straightening and splitting process. The spline of each ISV was manually traced for straightening, and the central plane of the z-stack was manually selected for splitting.

### Quantification of blood vessel diameter and length and EC number

To quantify vessel diameter and length, live *Tg(fli1:MYR-EGFP)^ncv^*^2^*, Tg(fli1:myr-mCherry)^ncv^*^1^, or *Tg(fli1:GAL4FF)^ubs^*^3^*, Tg(UAS:EGFP-UCHD)^ubs^*^18^ transgenic embryos were imaged at 2, 3, and 4 dpf. For EC number, live *Tg(fli1:h2bc1-mCherry)^ncv^*^31^ were embryos were imaged at the same stages. Confocal z-stacks were acquired using an Olympus UPLSAPO 40×/NA 1.25 silicone oil immersion objective. To determine ISV diameter, a line was drawn perpendicularly across the ISV in Fiji (NIH). The intensity profile along this line was used to identify two peaks corresponding to the vessel walls. A horizontal line was manually drawn at approximately half the maximum intensity of the peaks to measure the diameter. Three measurements were averaged for each ISV (No.10–18). The length of each ISV was measured by manually tracing a polyline along the vessel’s spine in Fiji. The tracing began at the point where the ISV connected to the DLAV and extended down to its connection with the DA or PCV. To count ECs, the nuclei within ISVs No.10– 18 were manually counted in Fiji.

### In vivo cell shape analysis

Single-cell labelling of ECs within ISVs was achieved through mosaic expression of different constructs: *fli1ep:lynEGFP* plasmid in wild-type embryos, *CxUAS:mylSbA2A3* for overexpression experiments, or *krit1* mutant cells in *Tg(fli1:GAL4FF)^ubs^*^3^*, Tg(UAS:EGFP-UCHD)^ubs^*^18^ transgenic background. At 2, 3, and 4 dpf, microangiography was performed, and vessels were imaged using an Olympus UPLSAPO 60×/NA 1.2 water immersion objective with an optical z-plane interval of 0.25 µm. Endothelial cell shape analysis was conducted semi-automatically using a custom-written Fiji script developed in Python, adapting methods from a previously described approach^13^. The analysis provided measures of cell area (A) and aspect ratio (R), which were subsequently utilized for strain analysis. Cell radial axis (C_r_) and vertical axis (C_v_) were calculated as: 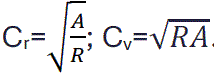. Effective cell number (E) was calculated as: 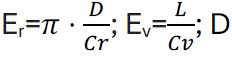, vessel diameter; L, vessel length. Vessel Strain (VS) was calculated as: 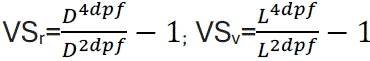; Cell strain (CS) was calculated as: 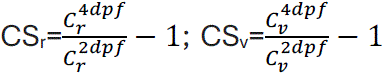. Effective strain due to cell number (ES) was calculated as: 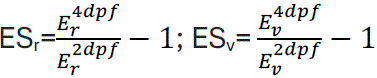.

### Analysis of actin organization

Processed images were analysed by splitting the blood vessel into front and back views to enable a clearer assessment of actin organization. Actin structure in the dorsal portion of the vessel, which was imaged as a representative region for the entire vessel (ISVs 10– 18), was classified manually as circumferential, mesh or longitudinal based on visible structural features. The percentage of organized actin was calculated from the total number of ISVs analysed across five independent experiments.

### Analysis of Actin and Myosin II Dynamics and Vessel Diameter

To analyse actin and myosin II dynamics, multichannel time-lapse z-stacks were maximally projected along the z-dimension to generate a 2D representation. A region of interest (ROI) covering the area of actin-myosin II colocalization was manually defined using a thick line. The average intensity profile for actin and myosin channels within the ROI was extracted at each time point and visualized as a kymograph. Intensity statistics for each time point were also extracted. Background intensity outside the vessel was also measured, and normalized intensities were calculated by dividing actin or myosin II signal by the background intensity. Vessel edges were traced from the kymograph, and the distance between the edges was measured to determine vessel diameter. Normalized actin and myosin II intensities, along with vessel diameter, were plotted over time to assess their dynamics and correlations.

### Laser ablation and Analysis of Mean Velocity of Recoiled Actin

Laser ablations were performed using a Zeiss LSM980 confocal microscope equipped with adjustable laser line MP-NDD (690–1300 nm) and a 63x Oil immersion objective (Plan-Apochromat, NA 1.2). Ablation parameters were controlled via ZEN (Blue 3.8) software. Each ablation consisted of a single laser pulse at 920nm with 80% laser power. Ablation was applied at the 5th frame of a time-lapse sequence comprising 50 frames captured at intervals of 250–300 ms (scan speed 13, and pixel time 0.75 µs), depending on the region of interest size. The location of ablation was manually drawn with a 0.3 µm × 6 µm box.

To quantify actin recoil velocity, the ablated region (a gap) in the actin network was analysed by drawing a line region of interest (ROI) across the gap to generate a kymograph. The edges of the gap were manually traced on the kymograph, and the distance between these edges was measured for each time frame to track the gap distance over time. Velocity was calculated by dividing the change in distance by the frame rate. The average velocity was calculated from measurements taken during the first 10 seconds post-ablation. For each ablation, three separate line ROIs were drawn across the gap, generating three independent average velocity measurements. The final mean velocity for each ISV was determined by averaging these three values. Each data point in the velocity bar chart represents the mean recoil velocity of an individual ablated ISV.

## Movie Legends

Supplementary Movie 1

(related to Fig. 2a). **Cell rearrangement and self-seam junction formation in aISV at 2-3 dpf**.

Time-lapse series of an aISV showing cortical and junctional actin in a *Tg(fli1:GAL4FF)^ubs^*^3^*; Tg(UAS:EGFP-UCHD)^ubs^*^18^ embryo at 2-3 dpf. The blood vessel is separated into front side (left channel) and back side (right channel). Timelapse started at 55 hpf and time interval is 30 mins. Time is in hours:minutes. Scale bar, 20 μm.

(related to supplementary Fig. 2a). **Cell rearrangement in aISV at 3-4 dpf**.

Time-lapse series of an aISV showing cortical and junctional actin in a *Tg(fli1:GAL4FF)^ubs^*^3^*; Tg(UAS:EGFP-UCHD)^ubs^*^18^ embryo at 3-4 dpf. The blood vessel is separated into front side (left channel) and back side (right channel). Timelapse started at 79 hpf and time interval is 30 mins. Time is in hours:minutes. Scale bar, 20 μm.

Supplementary Movie 3

(related to Fig. 2c). **Cell rearrangement and division in vISV at 2-3 dpf**.

Time-lapse series of a vISV showing cortical and junctional actin in a *Tg(fli1:GAL4FF)^ubs^*^3^*; Tg(UAS:EGFP-UCHD)^ubs^*^18^ embryo at 2-3 dpf. The blood vessel is separated into front side (left channel) and back side (right channel). Timelapse started at 55 hpf and time interval is 30 mins. Time is in hours:minutes. Scale bar, 20 μm.

Supplementary Movie 4

(related to supplementary Fig. 2c).

**Cell rearrangement and division in vISV at 3-4 dpf**.

Time-lapse series of a vISV showing cortical and junctional actin in a *Tg(fli1:GAL4FF)^ubs^*^3^*; Tg(UAS:EGFP-UCHD)^ubs^*^18^ embryo at 3-4 dpf. The blood vessel is separated into front side (left channel) and back side (right channel). Timelapse started at 79 hpf and time interval is 30 mins. Time is in hours:minutes. Scale bar, 20 μm.

Supplementary Movie 5

(related to Fig. 3c). **Circumferential actin bundles anchored to the cell membrane**.

Time-lapse series of a vISV segment showing circumferential actin anchoring to the cell membrane, in an embryo from *Tg(fli1:GAL4FF)^ubs^*^3^*; Tg(UAS:EGFP-UCHD)^ubs^*^18^ at 54 hpf. The right panel displays intensity-based image using Fire LUT in ImageJ, corresponding to the left panel. Time interval is 1 min. Time is in minutes:seconds. Scale bar, 5 μm.

Supplementary Movie 6

(related to Fig. 3d). **Circumferential actin bundles connected to cell-cell junctions**.

Time-lapse series of a vISV segment showing circumferential actin connecting to cell-cell junctions, in an embryo from *Tg(fli1:GAL4FF)^ubs^*^3^*; Tg(UAS:EGFP-UCHD)^ubs^*^18^ at 54 hpf. The right panel displays intensity-based image using Fire LUT in ImageJ, corresponding to the left panel. Time interval is 1 min. Time is in minutes:seconds. Scale bar, 5 μm.

Supplementary Movie 7

(related to Fig. 3d). **Circumferential actin bundles linking cell membrane and cell-cell junctions**.

Time-lapse series of a vISV segment showing circumferential actin bundles linked to both the cell membrane and cell-cell junctions, in an embryo from *Tg(fli1:GAL4FF)^ubs^*^3^*; Tg(UAS:EGFP-UCHD)^ubs^*^18^ at 54 hpf. The right panel displays intensity-based image using Fire LUT in ImageJ, corresponding to the left panel. Time interval is 1 min. Time is in minutes:seconds. Scale bar, 5 μm.

Supplementary Movie 8

(related to Fig. 4a). **Colocalization of circumferential actin bundle and myosin II**.

Time-lapse series of an aISV showing actin (EGFP-UCHD, middle panel) and non-muscle myosin II (myl9b-mCherry, right panel) in a *Tg(fli1:GAL4FF)^ubs^*^3^ *; Tg(UAS:EGFP-UCHD)^ubs^*^18^ and *Tg(CxUAS:mylSb-mCherry)^rk^*^32^ embryo at 52hpf. Colocalization of actin and myosin II at circumferential bundle is observed (arrows, merged channels at left panel). Time interval is 1 min. Time is in minutes:seconds. Scale bar, 5 μm.

Supplementary Movie 9

(related to Fig. 4b). **3D rendering of circumferential actomyosin bundle**.

3D rendering of an aISV showing colocalization of circumferential actin and myosin II (white arrows) at 8-minute from supplementary movie 8. 3D rendering is done using Huygens software. Myosin II expression is labelled in red and actin in green. Scale bar, 3 μm.

## Acknowledgements

We thank members of the Phng Lab for discussions and suggestions; Emi Taniguchi, RIKEN BDR Research Aquarium and RIKEN Kobe BioImaging Facilities & Factory for technical assistance; Hironobu Fujiwara for access to ZEISS LSM980 microscope; Holger Gerhardt and Anne Schmidt for plasmids; Heinz-Georg Belting and Markus Affolter for providing the transgenic line *Tg(fli1:GAL4FF)^ubs^*^3^, *Tg(UAS:EGFP-UCHD)^ubs^*^18^; and Fumio Motegi for comments on the manuscript. This work was supported by core funding from RIKEN BDR (to L-K.P.); RIKEN BDR DECODE project (to N.T.); RIKEN BDR STPJ Project (to L-K.P.); RIKEN BDR-Otsuka Pharmaceutical Collaboration Center (to Y.C.); the JSPS Grants-in-Aid for Scientific Research grants (22H022624 and 22H05168 to L-K.P; 22H05170 to S.O.; JP19H03394, JP19H05794, JP19H05795, JP22H02798, JP22H04926 to Y.O.); JST CREST grant JPMJCR1852 to Y.O.; JST Moonshot RCD grant JPMJMS2025-14 to Y.O.; AMED CREST grant JP23gm1700001s502 to Y.O.; ARC Discovery Project grant (DP230100393) to A.K.L.; NHMRC Ideas Grant (2029372) to A.K.L.; and LeDucq Transatlantic Network of Excellence “ReVAMP” (to L-K.P.).

## Contributions

L-K.P., Y.C. and N.T. conceptualized the project. Y.C., L-K.P., J.D.S., N.T., N.A., G.C., Y. O. and A.K.L. performed experiments. Y.C., N.T., L-K.P., I.K., N.A., G.C., and S.O. analysed data. Y.C. and N.T developed Fiji macros for image processing. Y.C. made the figures and movies. Y.C. and L-K.P. wrote and edited the manuscript. L-K.P. and A.K.L. acquired funding. L-K.P. supervised the project. All authors commented on the manuscript.

Correspondence to Li-Kun Phng.

## Ethics declarations

The authors declare no competing interests.

