## Supplemental Figures for "Circumferential actomyosin bundles drive endothelial cell deformations to constrict blood vessels"

**Supplementary Figure 1**

**a**, **b** Cell area and cell aspect ratio of aISV and vISV from 2 to 4 dpf. aECs, n=16/16//32 at 2/3/4 dpf; vECs, n=13/29/21 at 2/3/4 dpf. Statistical significance was assessed by ordinary one-way ANOVA with Turkey's multiple comparisons test. Mean values are indicated. **c** Average vessel area calculated from vessel diameter and length in aISV and vISV. aISV diameter n=228/284/213, length n=36/40/30 at 2/3/4 dpf; vISV diameter n=227/499/239, length n=39/59/33 at 2/3/4 dpf.

**Supplementary Figure 2**

**a** schematic shows EC exchange in aISV and vISV. **b** Number of cells that exchange between DLAV and aISVs and fractions of ISVs that are affected from 2-3 dpf (n=10 vessels) and 3-4 dpf (n=7 vessels). **c** Number of cells that exchange between DLAV and vISV, or between PCV and vISV, and fractions of ISVs that are affected from 2-3 dpf (n=5 vessels). **d**, **e** Number of mitotic events in ISVs and frequency of events from 2-3 dpf and 3-4 dpf.

**Supplementary Figure 3**

**a-c** Still images of an aISV (**a, b,** Supplementary Movie 3) or a vISV (**c,** Supplementary Movie 4) showing cortical and junctional actin in embryos from *Tg(fli1:GAL4FF)*$^{ubs3}$*; Tg(UAS:EGFP-UCHD)*$^{ubs18}$ at 3-4 dpf. The blood vessel is separated as the front side (left channel) and the back side (right channel). Schematics are traced from the original image and denote the constituent cells in colour (pink, blue, and yellow. Similar observations were made in 7 movies. Scale bar, 20um. **b** Magnification of the inset in **a**. Cross-section showed at z-plane indicated by black serrated line in **a**. Scale bar, 5um. **c** Cell divisions are observed and indicated by black arrows. Daughter cells (#1 and #2, or #3 and #4) are denoted in dark and light colours. Similar observations were made in 6 movies. Scale bar, 20um. ISV intersegmental vessel; aISV arterial ISV; vISV venous ISV.

**Supplementary Figure 4**

**a**, **b** Cell area and cell aspect ratio of aISV and vISV between control (myl9bA2A3-negative) and myl9bA2A3 overexpressing cells from 2 to 4 dpf. aECs, control, n=16/16//32, myl9bA2A3-OE, n=24/10/18 at 2/3/4 dpf; vECs, control, n=13/29/21, myl9bA2A3-OE n=12/22/27 at 2/3/4 dpf. Statistical significance was assessed by ordinary one-way ANOVA with Turkey's multiple comparisons test. Mean values are indicated.

**Supplementary Figure 5**

**a**, **b** Cell area and cell aspect ratio of aISV and vISV in transplanted *krit1* cells from 2 to 4 dpf. *krit1*$^{+/+}$, aEC n=4/11/13, vEC n=5/6/8 at 2/3/4 dpf), *krit1*$^{+/-}$, aEC n=17/12/14 , vEC n=10/14/20 at 2/4dpf) and *krit1*$^{-/-}$, aEC n=15/11/5 , vEC n=7/8/13 at 2/4 dpf). Statistical significance was assessed by ordinary one-way ANOVA with Turkey's multiple comparisons test. Mean values are indicated.

### Supplementary Figure 1

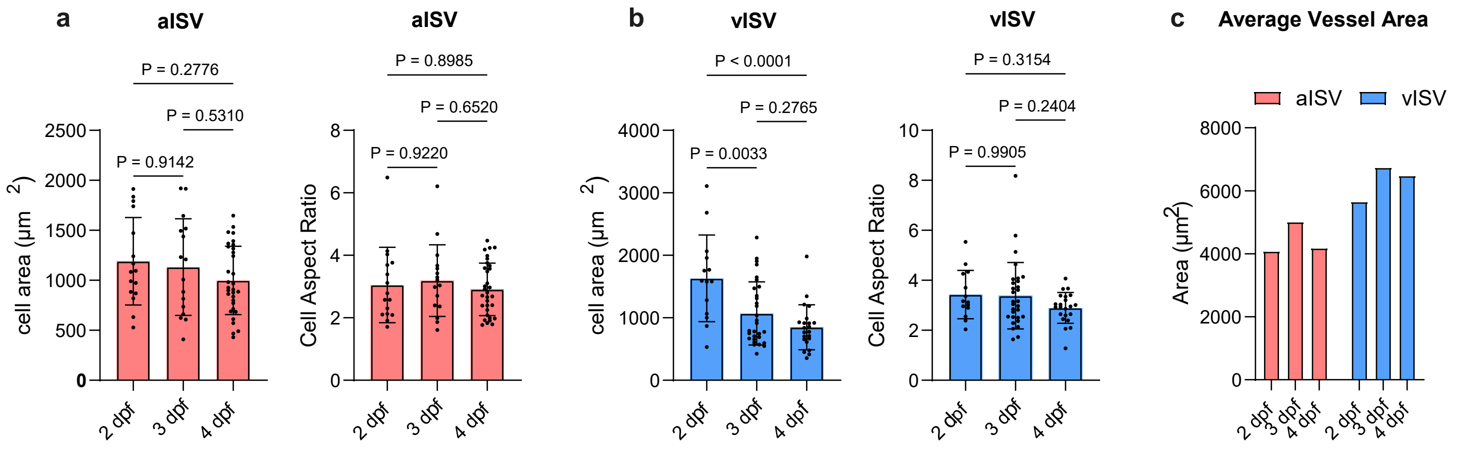

### Supplementary Figure 2

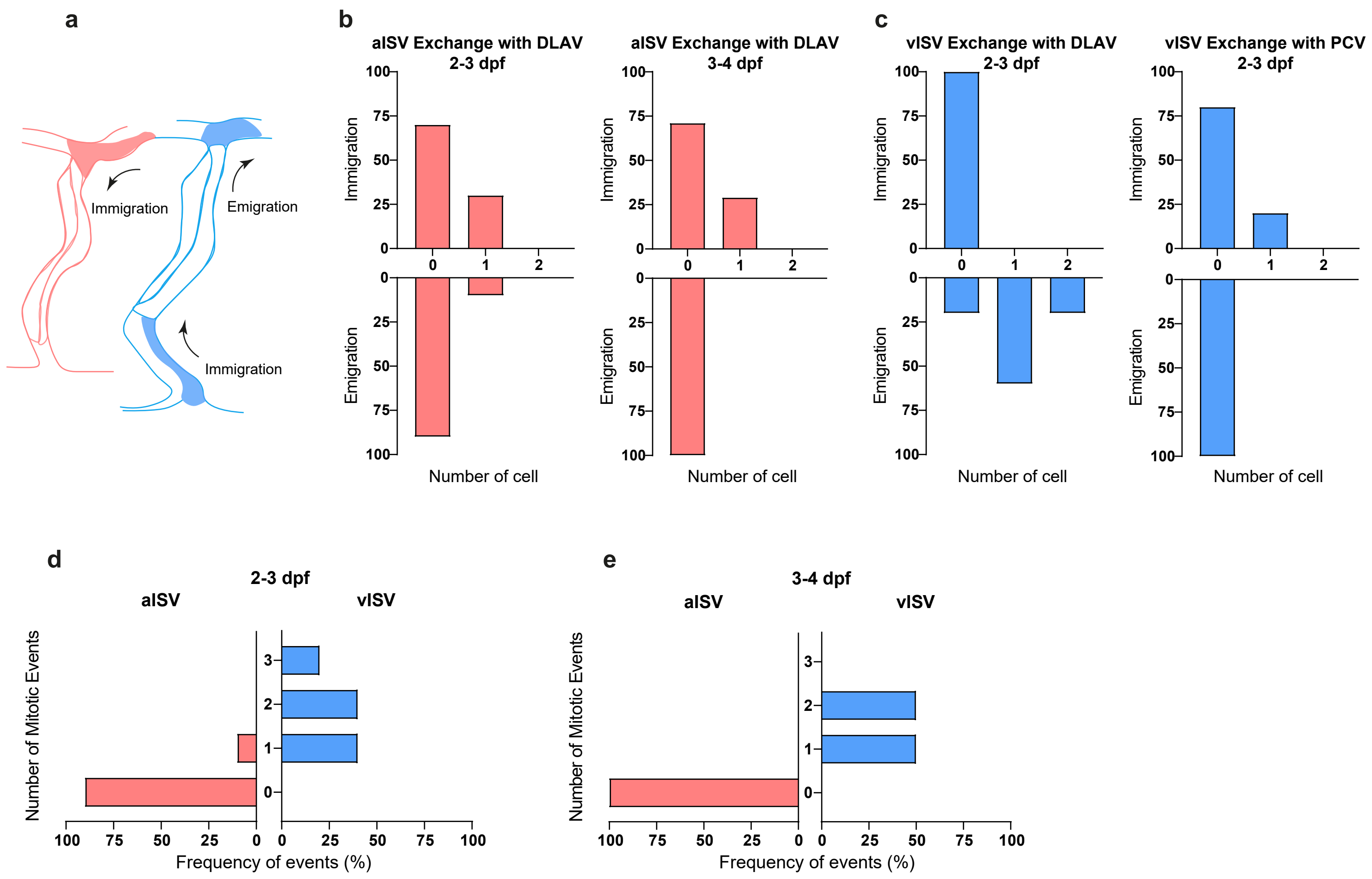

### Supplementary Figure 3

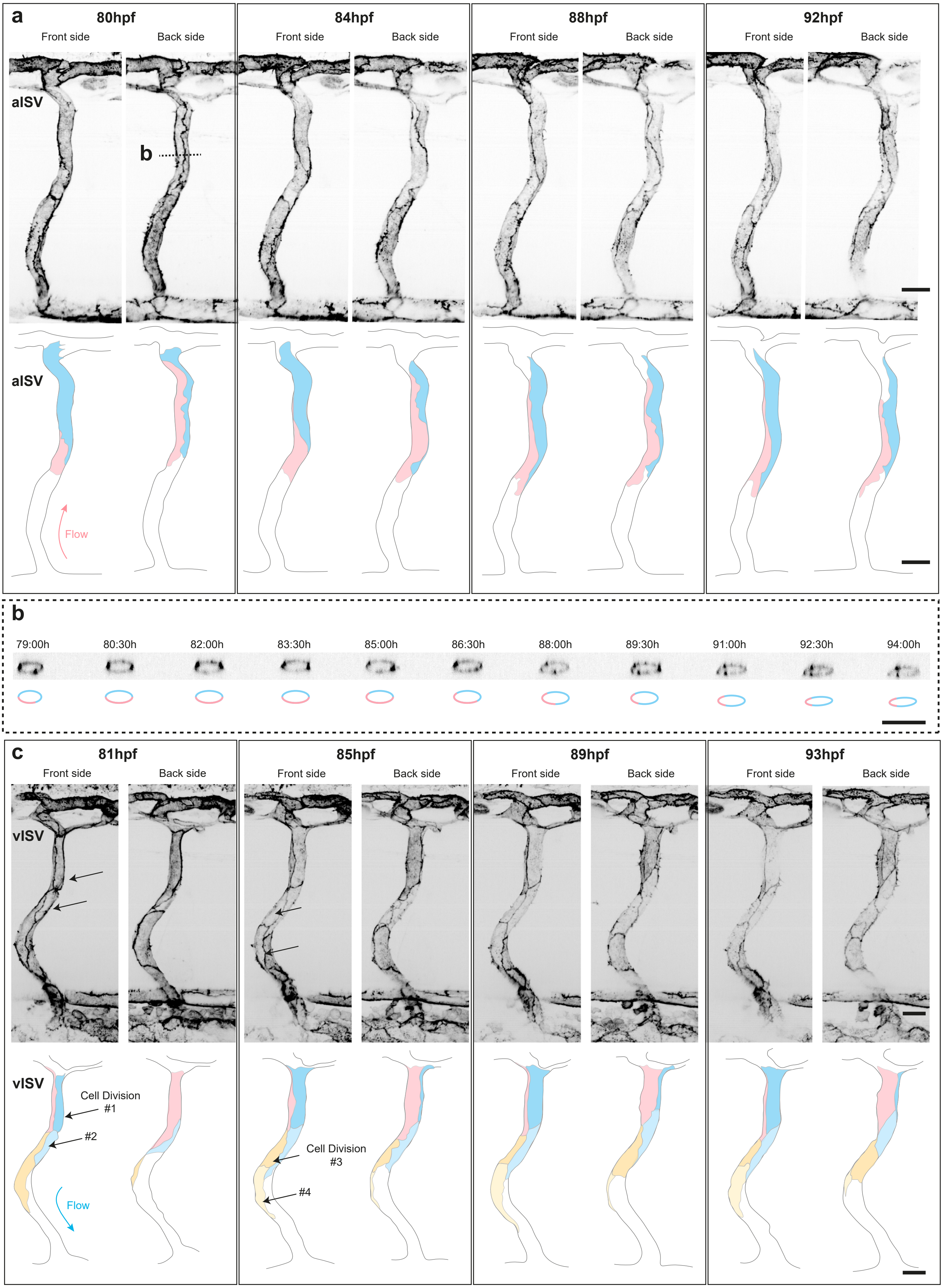

**Supplementary Figure 4**

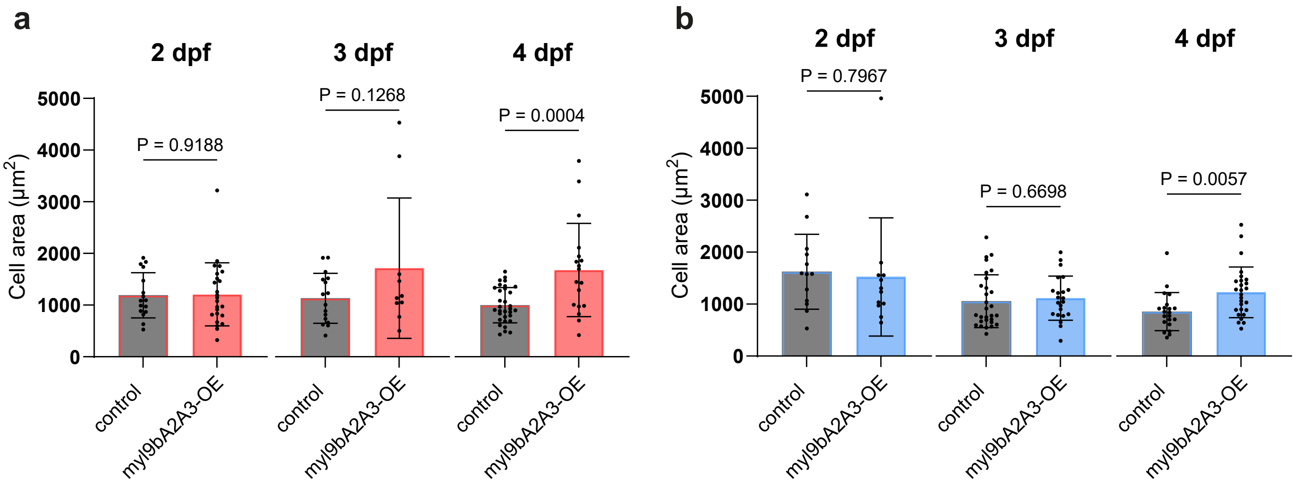

**Supplementary Figure 5**

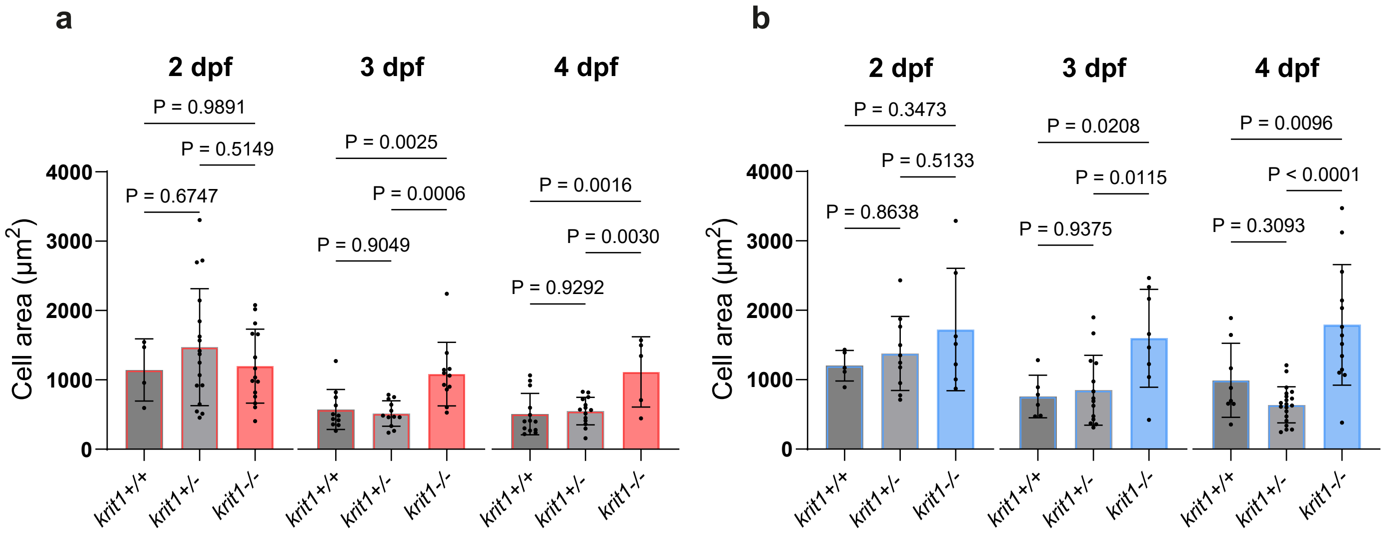
